# Bees differentiate sucrose solution from water at a distance

**DOI:** 10.1101/2022.05.20.492611

**Authors:** Melina Kienitz, Tomer Czaczkes, Massimo De Agrò

**Affiliations:** Animal Comparative Economics Lab, Department of Zoology, University of Regensburg, Germany; The BioRobotics Institute, Sant’Anna School of Advanced Studies, Pisa, Italy

**Keywords:** conditioning, vision, visual cues, cognition

## Abstract

Bees are perhaps the most important model for studying complex cognition in invertebrates, showing a variety of impressive abilities. Many experiments employ a training procedure in which the animal associates a characteristic of a “flower” (neutral stimulus) to sucrose solution (positive stimulus) over multiple foraging bouts. We hypothesized that sucrose solution might appear different from water to the bee sensorium, rendering it superfluous for the animal to learn the intended experimental cues. To test this hypothesis, we presented bumblebees simultaneously with 1.6M sucrose and water on artificial flowers. We presented the solutions in three conditions, to identify the most likely recognition mechanism: in drop form, inside cotton-plugged centrifuge tubes, and soaked into raised cigarette filters. The bumblebees chose to contact the sucrose solution significantly more often in all three conditions, confirming their ability to discriminate sucrose from water at a distance, with spectral differences being the most likely mechanism for this differentiation. These results have large implications for the design of training procedures, as the presence of an alternative cue may mask the learning of challenging tasks. This may force us to re-examine much of the literature and unpublished ‘negative results’ on bee cognition: weakening some claims, but strengthening others.

## Introduction

Western self-identity is strongly based on our own advanced cognitive abilities, as demonstrated even by the name we have given ourselves: *Homo sapiens*, the wise human. So too, our relationship to other animals is strongly influenced by our understanding of their own cognition, with ethical and legal protection provided to animal groups often depending on how cognitively advanced we consider them to be (Jones, 2013). Accurately identifying cognitive abilities thus has direct impacts on policy and consensus ethical opinion. Errors in both directions are costly: falsely identifying advanced cognition may have large economic impacts, while falsely rejecting advanced cognitive abilities may have large ethical repercussions. Invertebrates are currently the frontier of ethical consideration (Birch et al., 2021), especially as it becomes clear that brain organization may be more important than brain size in enabling cognitive abilities (Chittka and Niven, 2009). Indeed, a plethora of recent studies have emerged attesting the complexity of invertebrate behaviour and cognition (Cammaerts and Cammaerts, 2019; De Agrò et al., 2021, 2020; Finn et al., 2009; Howard et al., 2018; Jozet-Alves et al., 2013; Rößler et al., 2022)

Bees are probably the most important model organisms for studies on perception and cognition in invertebrates (Avarguès-Weber et al., 2012; Avarguès-Weber and Giurfa, 2013; Chittka and Waser, 1997; Gould, 1986; Howard et al., 2018; Menzel and Giurfa, 2001; Srinivasan, 2010; Zhang et al., 2010). Many such studies are made possible by the fact that these animals have evolved to be highly observant of their ever-changing environment (Baracchi, 2019), noticing and reacting to small differences in, for example, flower form, colour, illumination, and odour, to improve their foraging success (Dyer, 2006; Dyer et al., 2006; Dyer and Chittka, 2004; Lawson et al., 2017). Their ability to distinguish options increases when there is a combination of different cues acting together as multimodal signals, making bees more effective decision makers when flowers signal in more than one modality (Kulahci et al., 2008; Lawson et al., 2017).

When studying bee cognition, the use of conditioning techniques is very common. In some paradigms, the animals are trained to differentiate between ‘artificial flowers’ presenting opposing cues in one or more modalities. During training, the target flowers (Sn) can be presented in association with a reward (S+), usually consisting of sucrose solution (Biernaskie et al., 2009; Foster et al., 2014; Kulahci et al., 2008; Lotto and Chittka, 2005). The other flowers are associated with plain water or with a distasteful solution (S-), to maintain the same visual appearance between the options. The sucrose solution and the other unrewarding option are generally presented in the form of drops on the artificial flowers surface (Avarguès-Weber et al., 2010; Howard et al., 2018; Werner et al., 1988), or as centrifuge tubes containing each liquid, sometimes plugged with cotton, and placed in the artificial flowers (Avarguès-Weber et al., 2010; Cameron, 1981; Dyer, 2006; Foster et al., 2014).

However, surprisingly, the extent to which sucrose solution and water are distinguishable by the bees has been only sparsely investigated. Marden (1984) for example in one experiment tested three bumblebee foragers and found that they failed to distinguish sucrose from water if no supplementary cue was available (odour or spatial location). In that experiment the solutions were presented inside small tubes, a method analogous to the aforementioned use of centrifuge tubes. The author, however, did not test the bumblebees’ ability to discriminate drops placed on flowers’ surface. This is a critical omission, as these may provide more easily accessible information from multiple modalities. If bees can distinguish the reward and punishment directly, they may not attend to experiment-relevant cues. This becomes especially important in experimental setups that are cognitively challenging for bees. In such cases, the animals might resort to immediately available cues rather than attempting the intended task. We hypothesized a few possible sensory modalities through which bees, and specifically our model *Bombus terrestris*, may be able to discriminate solutions with and without sucrose at a distance.

The most likely candidate, in our opinion, is the visual modality. *Bombus terrestris* are trichromats, like most Hymenoptera, with blue, green and UV (ultraviolet) receptors (Menzel and Blakers, 1976). This means that bees perceive wavelengths between 300 and 600nm, and likely use this skill to identify UV-patterns on flowers (Chittka et al., 1994). This spectral sensitivity becomes crucial in the comparison between sucrose solution and water, as the two solutions are spectrally different over the whole range of visual spectrum of *B. terrestris*. Whether this difference is actually appreciable by *B. terrestris* or not has to be investigated (We provided the spectral signature of sucrose solution and water as supplement S1). Also note that quinine solution, a commonly-used negative stimulus, is also spectrally very different from sucrose solution in the UV range (See figure 8A in Sengar and Narula, 2019), and thus bees could differentiate the two solutions as well. Importantly, water and sucrose solution are not equally clear (dependent on the concentration of the sucrose solution). Even setting aside the solutions’ spectral signature, differential transparency may affect how the color of the surface in which the drop is placed is perceived (i.e. making it appear darker under the drop). We note that, on close observation, sucrose solution and water are visually distinct even to the human eye (see Figure 1, most apparent in 1C).

**Figure 1:**
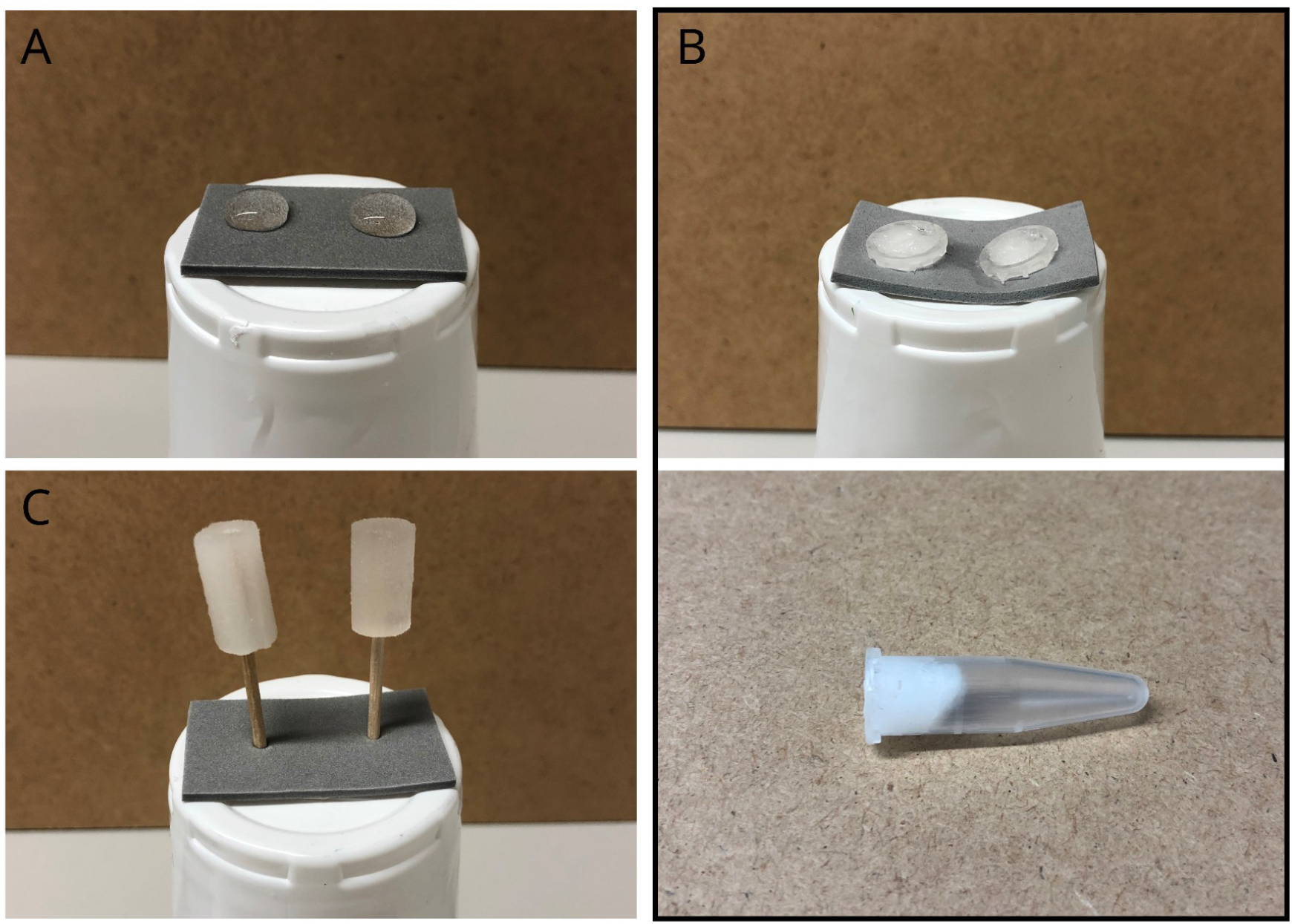
The three experimental conditions. In all of images, the right option is the sucrose solution, while the left is tap water. All were raised to 9cm on a plastic cup, and all were presented on a gray foam rubber rectangle. A) Condition 1: two drops were placed next to each using with a glass pipette. B) Condition 2: two centrifuge tubes were filled with cotton and then the cotton was soaked with one of the two options. The cotton is leveled at the rim of the tube. C) Condition 3: two soaked filters were affixed onto a toothpick and then placed next to each other on the artificial flower (note the visible difference between water and sucrose).

Remaining in the visual modality, bee compound eyes have a faster refresh rate than human eyes. Honeybees’ (*Apis mellifera*) critical flicker fusion frequency (CFF), the threshold above which a flickering light source start to be perceived as continuous, lies between 250 Hz and 265 Hz, four to five times higher than the human CFF (Ruck, 1958). Sucrose solution is also more viscose than water (Hidayanto et al., 2010) which leads to slower movement of the sucrose solution when shaken (SV1, or when gusts of air are produced, such as those generated by an approaching flying bee). When a bee flies to a drop of sucrose solution alongside a drop of water, the difference in viscosity might be visible for the bee, perhaps even more so than for humans due to the bees’ higher CFF rate (Hecht and Shlaer, 1936; Ruck, 1958).

Apart from vision, bees are also sensitive to chemical signals (Cameron, 1981; Lawson et al., 2017; Miyake et al., 1998) and like other nectar feeding animals (Von Arx et al., 2012), they can sense the humidity of flowers, and use this information to distinguish rewarding artificial flowers (Harrison and Rands, 2021). Sucrose solutions evaporate slower than water (Dittmar, 1935). Bees could therefore in principle sense the humidity difference between sucrose solutions and water, as sucrose solution would have lower local humidity.

It is often claimed that bees cannot smell sucrose solutions. However, support for this is poor and circumstantial. For example, when Avarguès-Weber et al. (2010) offered eight drops of sucrose solution and eight drops of quinine or water on top of a feeding platform the bees were not able to differentiate the drops by olfactory cues. However, rats can differentiate between different sucrose concentrations by olfactory cues alone, as measured the latency to lick time per concentration (Rhinehart-Doty et al., 1994). Bees may also be able to detect contaminants in sucrose, such as microorganisms growing in the solution, or their biproducts (i.e. volatile organic compounds). Honeybees detect microbial volatiles and can differentiate solutions containing microbiomes based on their volatiles (Rering et al., 2018), and bumblebees prefer feeding from yeast-inoculated artificial flowers (Schaeffer et al., 2017).

Other insects have been shown to visually discriminate solutions. The hoverfly *Eristalis arbustorum* can distinguish sucrose with or without 150ppb clothianidin at a distance, potentially due to their spectral difference (Clem et al., 2020). Unlike hoverflies, bees seem to have a non-gustatory preference for neonicotinoid-contaminated sucrose solutions, although the reason behind this preference and the mechanism is not clear (Bestea et al., 2021; de Brito Sanchez et al., 2005; Kessler et al., 2015).

Regardless of the modality employed, testing the ability of bees to discriminate between sucrose solution and water is a crucial step in insect cognition research. To this aim, we designed an experiment where bumblebees were allowed to forage in a flight arena in which we placed a grey artificial flower, simultaneously offering two solutions; 1.6M (548g/l) sucrose and tap water. Both solutions were sufficient to allow *at libitum* feeding. The bumblebee performed 9 consecutive foraging bouts in this setup, and for each bout we recorded the bee choice (i.e. which of the two options was contacted first), as well as if the animal inspected both solutions before choosing (i.e. hovered over both options before performing a decision, SV2). We hypothesize that if the bumblebees are able to discriminate sucrose solution from water they will choose the former significantly more often. We varied the presentation modality of the solution in order to narrow down the possible mechanisms enabling this discrimination. In a first condition, the two solutions were placed as large drops directly onto the flower surface (information available: visual difference, drop viscosity, evaporation speed, olfaction, Figure 1A). In a second condition, the solution was placed inside cotton-plugged centrifuge tubes (drop viscosity not appreciable, and requiring a steeper angle above the flower for visibility, Figure 1B). This should be similar to what Marden (1984) observed. In a third condition, the solutions were soaked into filter tips affixed to toothpicks (drop viscosity remains unavailable, but now the visibility is enhanced in respect to condition 2, Figure 1C). To ensure the validity of our “inspection” measure, we decided to video-record the bumblebees’ behaviour in the third condition and have a second, blinded, experimenter rescore the variable.

## Results

Results are summarized in Figure 2. Overall, the three conditions did not differ from each other (GLMM Analysis of Deviance, Chi-square=2.748, DF=2, p-value=0.253). We observed an effect of foraging bout number across all conditions (Chi-square=9.447, DF=1, p-value=0.002), as well as an effect of option inspection (whether the animals looked at both options before choosing, Chi-square=39.448, DF=1, p-value<0.0001)

**Figure 2:**
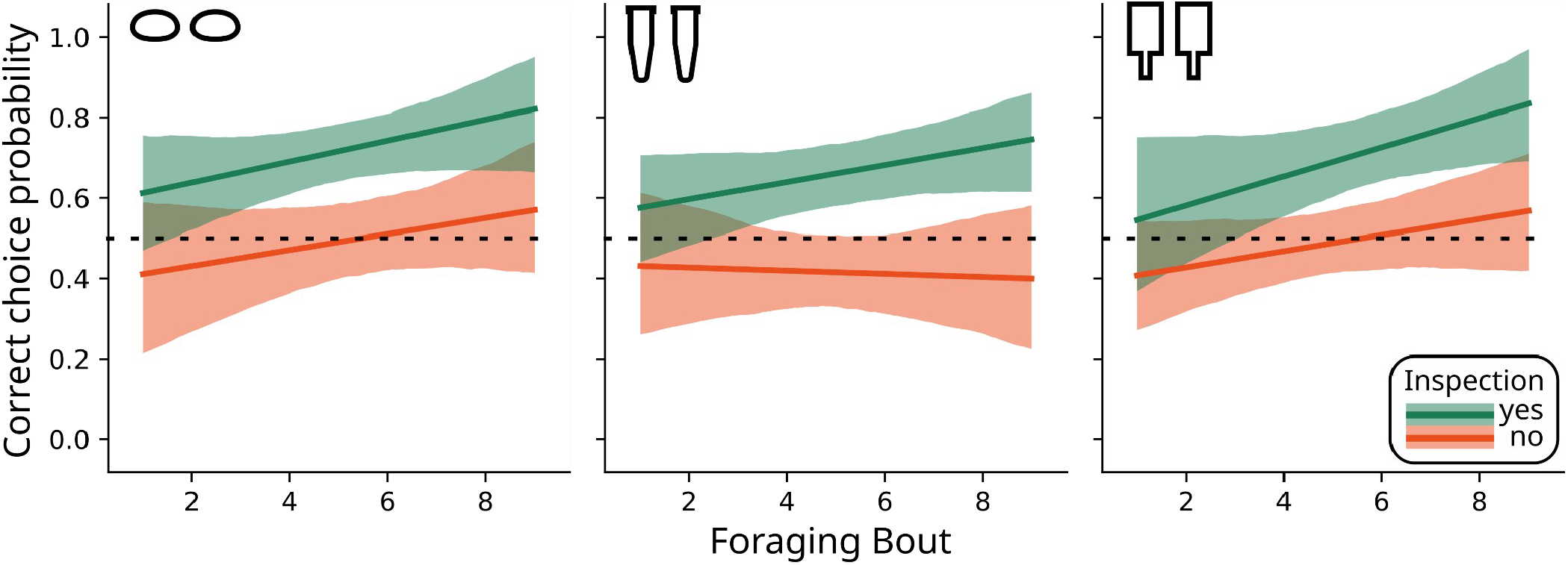
Results for the three experimental conditions. Y-axis represent probability of contacting first the sucrose solution. X-axis represents consecutive foraging bouts. Regression lines in green are for bees that inspected both stimuli before touching one in each bout (as scored live during the experiment). In red bees that touched directly one of them without looking at the other. Bees appeared overall capable of discriminate sucrose from water in the “drops” condition (left), in the “elevated” condition (right. Manually scored) but not in the “centrifuge tubes” one (center). In the latter, only bees that inspected both options appeared capable of discrimination.

Specifically, in the drop condition, bees were overall capable of discriminating the two solutions, choosing sucrose 60.5% of the times (post-hoc, Bonferroni corrected: estimate=0.4549, SE=0.133, t-ratio=3.41, p-value=0.0062). More specifically, bees that inspected both drops chose sucrose 72.1% of the time (estimate=0.9489, SE=0.185, t-ratio=5.118, p-value<0.0001), while the non-inspectors did so 49% of the times, a percentage not different from chance (estimate=-0.0391, SE=0.192, t-ratio=-0.204, p-value=1). We also found a non-significant trend towards increased choice accuracy, for both groups, across subsequent foraging bouts (inspected both options: trend=0.1296, SE=0.0736, t-ratio=1.762, p-value=0.4711; inspected one option: trend=0.08, SE=0.0732, t-ratio=1.093, p-value=1).

In the centrifuge tube condition, bees overall chose sucrose 53.9% of the times, a percentage not different from chance (post-hoc, Bonferroni corrected: estimate=0.1675, SE=0.128, t-ratio=1.304, p-value=1). When dividing the data into bees that inspected both options and bees that didn’t instead, we saw a significant preference for sucrose (66.3%) by the inspector group (estimate=0.6765, SE=0.176, t-ratio=3.832, p-value=0.0012). The animals that inspected only one option remained instead at a preference of 41.5%, not different from chance (estimate=-0.3415, SE=0.187, t-ratio=-1.827, p-value=0.6125). We again observed an upward, non-significant trend, for the bees that inspected both options, over subsequent visits (trend=0.0956, SE=0.0633, t-ratio=1.51, p-value=0.7893). The same trend was instead slightly downward when the bees inspected only one option (trend=-0.016, SE=0.0805, t-ratio=-0.201, p-value=1).

In the elevated feeder condition, bees chose sucrose 59.2% of the times, significantly more than chance level (post-hoc, Bonferroni corrected: estimate=0.3933, SE=0.135, t-ratio=2.907, p-value=0.0338). The group that inspected both options chose sucrose in 69.8% of visits (estimate=0.8368, SE=0.218, t-ratio=3.836, p-value=0.0012), while the other group remained at chance level, with a preference of 48.7% (estimate=-0.0501, SE=0.160, t-ratio=-0.313, p-value=1). We again observed an upward, non-significant trend, for the bees that inspected both options across visits (trend=0.1776, SE=0.086, t-ratio=2.065, p-value=0.2355) and for the bees that inspected only one (trend=0.0807, SE=0.0623, t-ratio=1.295, p-value=1).

Comparing the results of condition 3 when scored by hand vs using video recording, we found that the two experimenters were highly consistent in recognizing the bees first choice, with 88.8% of bouts being scored the same. Interestingly, for the remaining bouts, the videos revealed that bees sometimes quickly extended their proboscis and tasted one of the options without landing, at a speed unnoticeable during real-time scoring. Regarding the inspection behaviour, only 58% of trials were scored equally. this percentage is, frankly, insufficient for inspection to be considered as a solid measure, even for the other conditions, as it seems to be too dependent on the experimenter judgment. As such, we advise caution for interpretations linked to this behaviour, while we are highly confident about the overall choice pattern. We followed up by analyzing the video-scored data using the same procedure as before, in order to be sure that the results would take the same direction even when considering the fast rapid tasting behaviour (and so revealing a different choice). Bees chose sucrose 64.3% of the times, significantly more than chance level (post-hoc, Bonferroni corrected: estimate=0.618, SE=0.163, t-ratio=3.799, p-value=0.0005). The group that inspected both options chose sucrose in 74.6% of visits (estimate=1.075, SE=0.290, t-ratio=3.707, p-value=0.0008), while the other group remained at chance level, with a preference of 54%% (estimate=-0.148, SE=0.148, t-ratio=1.093, p-value=0.8263). We again observed an upward, non-significant trend, for the bees that inspected both options across visits (trend=0.2627, SE=0.121, t-ratio=2.176, p-value=0.061) and for the bees that inspected only one (trend=0.0934, SE=0.057, t-ratio=1.64, p-value=0.2046).

## Discussion

In this experiment, we found that bumblebees were able to discriminate sucrose from water solution in conditions 1 and 3. The results of condition 2 (centrifuge tubes) remains unclear, as the presence of the effects depends on the inclusion or exclusion of bees that did not inspect both options, a difficult behavior to score consistently. Given the consistent difference between inspected and non-inspected visits, we tend to believe that the ‘inspection’ variable does capture some real differences between the inspection and non-inspection visits. Given the consistency of condition 2 with the results from Marden (1984), it seems that indeed hiding the drop inside a tube is an effective solution to prevent solutions discrimination.

From our experiment alone, it is impossible to unequivocally identify which mechanisms and information bees use to discriminate the two solutions. However, given the results, some mechanisms seem more likely than others.

It is likely that the solution viscosity, and consequent vibration speed of the drops, has little or no importance for the task. In fact, bumblebees were equally successful in condition 3 (elevated), where the liquid was soaked into a filter and the “drops” could not shake. It is less clear whether olfaction or hygroscopy is being used. If we were to consider condition 2 as unsuccessful, and that the bees failed to distinguish the two solutions in this treatment, the use of chemical cues has to be excluded. Indeed, in all three conditions the presence of volatiles should be similar. However, even if the ‘inspection’ variable is to be trusted, there should be no reason why the lack or presence of hovering should have a different effect on the three conditions: either hovering over the rewards is required to sense volatiles in all conditions, or it is never required. We thus believe olfaction or hygroscopy is unlikely to be the differentiation mechanism.

In our opinion, vision remains the most likely mechanism. It is unclear whether the spectral difference between the drops is perceptible to the bumblebees, or whether they use other physical properties related to light-matter interactions, such as differential light scattering of the surface below. Certainly the drops appear differently “coloured” in all three conditions to the human eye (most noticeably in the third condition, see Figure 1C). The ability to perceive such difference visually by humans is not directly extendable to the bees *sensorium*, however our results suggest that they do. In conditions 1 and 3, the bumblebees seem to improve across trials even without inspection, which can be explained by the ease with which the visual information can be acquired from afar during flight. Condition 2 (centrifuge tubes) instead partially hides the visual information, so that it can only be accessed from a steep angle above the artificial flowers.

Regardless of the mechanism behind this discrimination ability, its mere presence has wide-reaching implications for researchers studying insect cognition and behavioural ecology. Many dual-choice experiments performed on bees employ the precaution of completely removing the reward during crucial test trials, or use stimuli presentation that precede reward availability, rather than presenting them concurrently. As such, we feel that this new finding does not jeopardize many positive observations made by previous experiments. On the other hand, especially experiments where positive and negative stimuli are present in test visits, and experiments where training shows a higher success rate than the unrewarded tests, might be consistently underestimating effects due to bees relying directly on the visual properties of sucrose. For example, in a recent series of studies by Howard et al. (2019a, 2019b, 2019c, 2018) examining honeybees’ numerical cognition abilities, honeybees were trained on drops of sucrose below the relevant stimuli. The authors report consistently higher effect sizes in the training phase than in the test and transfer phases (where reward solution was not available). It is possible that some individuals learned to discriminate the solutions themselves, rather than the training stimuli, and were unable to perform once these were removed. Similar concerns can be raised for other studies examining cognition in bees (e.g. Avarguès-Weber et al., 2011; Muszynski and Couvillon, 2015).

It is unclear at this point to what extent the ability to distinguish sucrose from water at a distance has impacted past results. It is likely that simple discrimination tasks using highly salient cues (e.g. flower colours) are unaffected. Less salient cues, such as those with subtle differences used in studies of advanced cognition, are likely more strongly affected. Specific procedural details, such as the reward volume or concentration, or the flower colour and material, may impact the bees’ perception, partially masking this effect. Some experiments indeed employ procedures which may hide visual sucrose cues: it has been suggested that bees cannot see inside centrifuge tubes at a certain angle (Giurfa et al., 1996), which seem to be consistent with the results we observed in condition 2. However, we could not definitely exclude the availability of non-visual cues, nor that centrifuge tubes completely hide visual cues. Fully quantifying the influence of this sucrose discrimination ability on past experiments would require a comparison of every experimental cue with every sucrose reward presentation to define relative relevance – something beyond the scope of this work. In the absence of these direct comparisons, the visuo-chemical characteristics of the options should be considered a relevant cue, until demonstrated otherwise. It is for example possible that drops <2μl are too small to show a difference, but this would need to be directly tested, rather than assumed.

The ability to discriminate sucrose from water at a distance may have prevented the success of previous procedures. This is hard to quantify, since due to a positive publication bias unsuccessful experiments often remain unreported (Duyx et al., 2017; Mlinarić et al., 2017). This is especially unfortunate, as we may be missing out on surprising skills and abilities that these insects are capable of, research into which has been abandoned due to this unforeseen confounding factor. On an important and positive note, we may have been *underestimating* the ease with which bees can learn abstract concepts, suggesting that previous studies demonstrating the advanced cognitive powers of bees (e.g. Avarguès-Weber et al., 2011; Howard et al., 2018) are even more ecologically relevant. Overall, we hope that our findings will inspire a reanalysis of the literature with a critical eye, both for unexpected positive and negative results. Moreover, we hope that our findings will encourage researchers to re-examine their past work and previously abandoned ideas, which may give rise to new and unexpected discoveries.

## Materials and methods

### Subjects

We used 6 commercial *Bombus terrestris* colonies, some queenright and some queenless (Biological systems, Naturpol). Bumblebees had continuous access to a flight arena (60×45×28 cm), covered with a transparent Plexiglas top. The rest of the arena was constructed of grey plastic and wood. Two transparent doors (16×16cm) provided hand access. The bottom of the flight arena was covered with standard printer paper. For testing purposes, six equidistant dots were drawn on the layer of paper, which were later used for determining the flower locations. The flight arena and nest box were connected via a transparent tunnel with sliding doors to control access. From Friday afternoon until Monday morning the bumblebees were given access to *ad libitum* 1M sucrose solution via a feeder in the flight arena, supplemented with pollen provided twice per week directly into the nest. Feeders were elevated to 9 cm, to accustom bumblebees to forage at this height. During testing time, food was only provided through the training procedure. A total of 90 individuals, 30 per condition, were tested. The colonies were kept on a 14.5:9.5 light:dark cycle. Light-conditions in the flight arena (Supplement S1) were measured with a Compact Spectrometer (Thorlabs CCS200/M, Extended Range: 200 – 1000 nm). The room was lit with ceiling mounted CFL light tubes and additional LED lights nearer the flight arena.

### Experimental procedure

#### Pre-training

Pre-training took place from Monday-Thursday, training and testing took place Tuesday-Friday. Before testing, a colony had to be pre-trained for at least one day, in order to identify active foragers and to teach them how to feed from the artificial flowers. The artificial flowers that were used for this experiment consisted of a grey foam rubber rectangle, placed on an upside-down white plastic cup (elevation 9 cm). 1M sucrose solution made with tap water was placed in the centre of the artificial flower. Depending on the experiment, it was either presented in one of three different feeder types: as a single large drop (drop-feeder), in a centrifuge tube filled with soaked cotton (centrifuge-tube feeder) or as a soaked cigarette filter (8 mm) stuck on a toothpick (elevated-feeder). The differently shaped feeders corresponded with the three experimental conditions. Pre-training feeders were placed along the vertical axis of the flight arena, avoiding the six testing positions.

Bees were pre-trained to use the feeders for at least one day prior to testing, by offering the feeder type tested (see below) with no water alternative. Pre-training allowed active foragers to be identified and marked prior to testing.

#### Testing

While feeding from the pre-training artificial flower, bees were marked with acrylic paint on the top of the thorax, abdomen or legs using a toothpick. If the marked bee then carried out two consecutive foraging bouts, it proceeded to the experiment.

An artificial flower was placed in the area (see below for details). The flower offered two options, no more than 15 mm from each other. The rewarding option was 1.6M sucrose solution made with tap water, the unrewarding option was tap water. Presentation method depending on the experiment: two drops (drop-feeders), centrifuge tubes filled with soaked cotton (centrifuge-tube-feeders), or soaked cigarette filters stuck on a toothpick (elevated-feeders, Figure 1). The flower was randomly placed in one of six positions. The relative position of the sucrose to the water option was in one of four possible positions (1= right, 2= left, 3= front, 4= back; oriented as if entering the flight arena). The bumblebees were therefore not able to learn or predict the position of the flower, or the position of the sucrose relative to the water.

To begin a bout a single marked bumblebee was allowed into the flight arena and allowed to choose between the two options (1.6M sucrose solution and water) on the artificial flower. It then consumed the sucrose solution until satiation and was let back into the nest. The artificial flower was then removed from the arena, and substituted with a clean one. Each bumblebee performed nine foraging bouts, and with every new bout the positions of the artificial flower, and relative positions of sucrose solution and water, were randomized. For each bout, we recorded the positions of the flower and sucrose option, the first choice between the options, the time from entering the arena to first contacting the flower), whether the bumblebee rejected the solution it chose first, which option the bumblebee chose to feed from, if the bumblebee drank to satiation, and – importantly – whether the bumblebee inspected both options or just one option before making its first choice, and if only one option, which option was inspected. We defined the first choice as a bumblebee coming in contact with one of the presented options. We defined inspected both options as either a) when a bumblebee hovered facing both options right before choosing one, b) when a bumblebee hovered and faced one option and then chose the other option, or c) when a bumblebee landed on the platform without touching the options and then chose one (see video SV2).

Experiments were repeated using three different sucrose presentation methods (Figure 1). In condition 1 we presented the bumblebees with two equally-sized drops onto the same artificial flower (Figure 1A). Artificial flowers were grey foam rubber rectangles elevated to 9 cm on plastic cups. The aim of this experiment was to test whether foragers of *B. terrestris* can distinguish a drop of 1.6M sucrose solution from a drop of water without touching and tasting the cue first.

After having observed that bees were successful in discriminating the two drops in condition 1 (see results), we designed a second condition. Since 1.6M sucrose solution is more viscose than water, and this difference is noticeable to a human observer, as the drops vibrate differently when blown on or moved (see video SV1). The aim of this second condition was to remove the viscosity differences of the cues. This was achieved by presenting the sucrose solution and water in cotton-filled 1.5ml centrifuge tubes, which pierced the grey foam rectangle so that their lip was level with the foam (Figure 1B). Every test tube was used once, and the lid was cut off. Every tube was filled with a wad of cotton and soaked with a pipette. The cotton was pushed down into the tube to a similar level for both the sucrose and water treatments. The cotton was thus only visible from above the feeders (minimum circa 30 degrees from vertical).

A third condition was run to further explore the potential cues bumblebees might be using to do this. Here, we enhanced the visibility of the two solutions, while simultaneously obscuring any structural cues. This was achieved by soaking cigarette filters and affixing them onto toothpicks. These elevated feeders where thus very evident (and looked different at this point even to the human observer, Figure 1C), but not being presented in drop form removed structural information.

#### Video analysis

To ensure consistency in the scoring procedure, video recordings of condition 3 was also scored by a blind experimenter. In the video analysis we recorded the bee’s first choice of options, if it rejected the first choice the option that the bumblebee finally chose to feed from, what option the bumblebee inspected before deciding, and if the bumblebee tasted an option instead of visually distinguishing it. This last behaviour was noticed during initial video inspection: occasionally, bumblebees would rapidly flick their proboscis out to the feeder and lick it before making a choice. This behaviour is extremely rapid, and almost impossible to see in real-time.

When bumblebee tasted the option instead of inspecting it from a distance, this was considered as the first choice. Some bumblebees also inspected both options prior to performing a tasting. Also, for these bumblebees, we considered the option it tasted as its first choice.

#### Spectral measurements

To test whether water and 1.6M sucrose solution differ in their spectral absorbance, we carried UV-Vis spectroscopic measurements (ultraviolet-visible-spectroscopy) on both solutions using a Varian Cary 50 Bio and 10 × 10 mm cuvettes. We also measured the absorbance, transmittance, and reflectance of sucrose solution with tap water as the baseline. The sucrose solution was made using LC-MS Grade water. Comparing the absorbance spectrum to the visible spectrum of *B. terrestris* (Skorupski et al., 2007) allows us to ascertain whether the sucrose solution and water could in principle be visually different for the bee.

#### Statistical Analysis

The entire statistical analysis, including figure code and analysis results, is presented in supplement S2. Raw data is available in supplement S1. All the statistical analyses were performed in R 4.1.2 (R Core Team, 2020). We initially checked the effect of incidental measurements, namely the effect of the randomized positioning of rewards on overall preference and the duration of every foraging bout. The results for these two measures are not reported here, as they had no effect of interest. They are reported in the supplement for completeness.

For our main analysis, we observed the probability of choosing sucrose solution, across foraging bouts and depending on whether the animals inspected both options. We employed a generalized mixed effect model with a binomial distribution using the package glmmTMB (Brooks et al., 2017; Magnusson et al., 2020). The goodness of fit was evaluated with the package “DHARMa” (Hartig, 2018). We performed an analysis of deviance to observe the effect of the predictors using the package car (Fox and Weisberg, 2019), and then performed Bonferroni-corrected post-hoc analysis on predictors that have an effect using the package emmeans (Lenth, 2018).

We also performed pearsons correlation tests to check the consistency between manually scored data and video scoring for condition 3. This was performed both on drop choice and on the inspection variable.

After analysis, the data was then passed onto a Python 3 (Van Rossum and Drake, 2009) environment using the package reticulate (Ushey et al., 2021), to produce graphs. To achieve this, we used the libraries pandas (Jeff Reback et al., 2020), numpy (Oliphant, 2006; van der Walt et al., 2011), matplotlib (Hunter, 2007) and seaborn (Waskom et al., 2017).

## Supporting information

SI - raw data

SI - analysis script

Video 1 - drop viscosity

Video 2 - inspection of drop

## Acknowledgments

We would like to thank Professor David Baracchi for input and suggestions provided on the manuscript. MDA was funded through a European Research Council Starter Grant (938181) to TJC. TJC was funded through a Heisenberg fellowship from the Deutsche Forschungsgemeinschaft (CZ 237 / 4-1).

## References

Avarguès-Weber A, Dyer AG, Combe M, Giurfa M. 2012. Simultaneous mastering of two abstract concepts by the miniature brain of bees. PNAS 109:7481–7486. doi:10.1073/pnas.1202576109

Avarguès-Weber A, Dyer AG, Giurfa M. 2011. Conceptualization of above and below relationships by an insect. Proceedings of the Royal Society B: Biological Sciences 278:898–905. doi:10.1098/rspb.2010.1891

Avarguès-Weber A, Giurfa M. 2013. Conceptual learning by miniature brains. Proc R Soc B 280:20131907. doi:10.1098/rspb.2013.1907

Avarguès-Weber A, Sanchez MG de B, Giurfa M, Dyer AG. 2010. Aversive Reinforcement Improves Visual Discrimination Learning in Free-Flying Honeybees. PLOS ONE 5:e15370. doi:10.1371/journal.pone.0015370

Baracchi D. 2019. Cognitive ecology of pollinators and the main determinants of foraging plasticity. Current Zoology 65:421–424. doi:10.1093/cz/zoz036

Bestea L, Réjaud A, Sandoz J-C, Carcaud J, Giurfa M, de Brito Sanchez MG. 2021. Peripheral taste detection in honey bees: What do taste receptors respond to? Eur J Neurosci 54:4417–4444. doi:10.1111/ejn.15265

Biernaskie JM, Walker SC, Gegear RJ. 2009. Bumblebees Learn to Forage like Bayesians. The American Naturalist 174:413–423. doi:10.1086/603629

Birch J, Burn C, Schnell A, Browning H, Crump A. 2021. Review of the Evidence of Sentience in Cephalopod Molluscs and Decapod Crustaceans. Animal Feeling - General.

Brooks ME, Kristensen K, Benthem KJ van, Magnusson A, Berg CW, Nielsen A, Skaug HJ, Maechler M, Bolker BM. 2017. glmmTMB Balances Speed and Flexibility Among Packages for Zeroinflated Generalized Linear Mixed Modeling. The R Journal 9:378–400.

Cameron SA. 1981. Chemical signals in bumble bee foraging. Behav Ecol Sociobiol 9:257–260. doi:10.1007/BF00299880

Cammaerts M-C, Cammaerts R. 2019. Ants’ notion of zero through the perception of the absence of an odor. International Journal of Biology 11:1–12.

Chittka L, Niven J. 2009. Are Bigger Brains Better? Current Biology 19:R995–R1008. doi:10.1016/j.cub.2009.08.023

Chittka L, Shmida A, Troje N, Menzel R. 1994. Ultraviolet as a component of flower reflections, and the colour perception of hymenoptera. Vision Research, The Biology of Ultraviolet Reception 34:1489–1508. doi:10.1016/0042-6989(94)90151-1

Chittka L, Waser NM. 1997. Why Red Flowers Are Not Invisible to Bees. Israel Journal of Plant Sciences 45:169–183. doi:10.1080/07929978.1997.10676682

Clem S, Sparbanie T, Luro A, Harmon-Threatt A. 2020. Can anthophilous hover flies (Diptera: Syrphidae) discriminate neonicotinoid insecticides in sucrose solution? PLOS ONE 15:e0234820. doi:10.1371/journal.pone.0234820

De Agrò M, Oberhauser FB, Loconsole M, Galli G, Dal Cin F, Moretto E, Regolin L. 2020. Multimodal cue integration in the black garden ant. Anim Cogn 23:1119–1127. doi:10.1007/s10071-020-01360-9

De Agrò M, Rößler DC, Kim K, Shamble PS. 2021. Perception of biological motion by jumping spiders. PLOS Biology 19:e3001172. doi:10.1371/journal.pbio.3001172

de Brito Sanchez MG, Giurfa M, de Paula Mota TR, Gauthier M. 2005. Electrophysiological and behavioural characterization of gustatory responses to antennal “bitter” taste in honeybees. Eur J Neurosci 22:3161–3170. doi:10.1111/j.1460-9568.2005.04516.x

Dittmar JH. 1935. Hygroscopicity of Sugars and Sugar Mixtures. Ind Eng Chem 27:333–335. doi:10.1021/ie50303a021

Duyx B, Urlings MJE, Swaen GMH, Bouter LM, Zeegers MP. 2017. Scientific citations favor positive results: a systematic review and meta-analysis. Journal of Clinical Epidemiology 88:92–101. doi:10.1016/j.jclinepi.2017.06.002

Dyer AG. 2006. Bumblebees directly perceive variations in the spectral quality of illumination. J Comp Physiol A 192:333. doi:10.1007/s00359-005-0088-z

Dyer AG, Chittka L. 2004. Bumblebee search time without ultraviolet light. Journal of Experimental Biology 207:1683–1688. doi:10.1242/jeb.00941

Dyer AG, Whitney HM, Arnold SEJ, Glover BJ, Chittka L. 2006. Behavioural ecology: Bees associate warmth with floral colour. Nature 442:525. doi:10.1038/442525a

Finn JK, Tregenza T, Norman MD. 2009. Defensive tool use in a coconut-carrying octopus. Current Biology 19:R1069–R1070. doi:10.1016/j.cub.2009.10.052

Foster JJ, Sharkey CR, Gaworska AVA, Roberts NW, Whitney HM, Partridge JC. 2014. Bumblebees Learn Polarization Patterns. Current Biology 24:1415–1420. doi:10.1016/j.cub.2014.05.007

Fox J, Weisberg S. 2019. An R Companion to Applied Regression, Third. ed. Thousand Oaks CA: Sage.

Giurfa M, Vorobyev M, Kevan P, Menzel R. 1996. Detection of coloured stimuli by honeybees: minimum visual angles and receptor specific contrasts. J Comp Physiol A 178:699–709. doi:10.1007/BF00227381

Gould JL. 1986. Pattern learning by honey bees. Animal Behaviour 34:990–997. doi:10.1016/S0003-3472(86)80157-9

Harrison AS, Rands SA. 2021. The ability of bumblebees Bombus terrestris (Hymenoptera: Apidae) to detect floral humidity is dependent upon environmental humidity 2021.08.13.456254. doi:10.1101/2021.08.13.456254

Hartig F. 2018. DHARMa: Residual Diagnostics for Hierarchical (Multi-Level / Mixed) Regression Models.

Hecht S, Shlaer S. 1936. Intermittent stimulation by light: the relation between intensity and critical frequency for different parts of the spectum. Journal of General Physiology 19:965–977. doi:10.1085/jgp.19.6.965

Hidayanto E, Tanabe T, Kawai J. 2010. Measurement of viscosity and sucrose concentration in aqueous solution using portable Brix meter. Berkala Fisika 13:23–28.

Howard SR, Avarguès-Weber A, Garcia JE, Greentree AD, Dyer AG. 2019a. Surpassing the subitizing threshold: appetitive–aversive conditioning improves discrimination of numerosities in honeybees. Journal of Experimental Biology 222:jeb205658. doi:10.1242/jeb.205658

Howard SR, Avarguès-Weber A, Garcia JE, Greentree AD, Dyer AG. 2019b. Symbolic representation of numerosity by honeybees (Apis mellifera): matching characters to small quantities. Proceedings of the Royal Society B: Biological Sciences 286:20190238. doi:10.1098/rspb.2019.0238

Howard SR, Avarguès-Weber A, Garcia JE, Greentree AD, Dyer AG. 2019c. Numerical cognition in honeybees enables addition and subtraction. Science Advances 5:eaav0961. doi:10.1126/sciadv.aav0961

Howard SR, Avarguès-Weber A, Garcia JE, Greentree AD, Dyer AG. 2018. Numerical ordering of zero in honey bees. Science 360:1124–1126. doi:10.1126/science.aar4975

Hunter JD. 2007. Matplotlib: A 2D graphics environment. Computing in Science & Engineering 9:90–95. doi:10.1109/MCSE.2007.55

Jeff Reback, Wes McKinney jbrockmendel, Joris Van den Bossche, Tom Augspurger, Phillip Cloud, gfyoung, Sinhrks, Adam Klein, Matthew Roeschke, Simon Hawkins, Jeff Tratner, Chang She, William Ayd, Terji Petersen, Marc Garcia, Jeremy Schendel, Andy Hayden MomIsBestFriend, Vytautas Jancauskas, Pietro Battiston, Skipper Seabold, chris-b1, h-vetinari, Stephan Hoyer, Wouter Overmeire alimcmaster1, Kaiqi Dong, Christopher Whelan, Mortada Mehyar. 2020. pandas-dev/pandas: Pandas 1.0.3. Zenodo. doi:10.5281/zenodo.3715232

Jones RC. 2013. Science, sentience, and animal welfare. Biol Philos 28:1–30. doi:10.1007/s10539-012-9351-1

Jozet-Alves C, Bertin M, Clayton NS. 2013. Evidence of episodic-like memory in cuttlefish. Curr Biol 23:R1033–1035. doi:10.1016/j.cub.2013.10.021

Kessler SC, Tiedeken EJ, Simcock KL, Derveau S, Mitchell J, Softley S, Radcliffe A, Stout JC, Wright GA. 2015. Bees prefer foods containing neonicotinoid pesticides. Nature 521:74–76. doi:10.1038/nature14414

Kulahci IG, Dornhaus A, Papaj DR. 2008. Multimodal signals enhance decision making in foraging bumble-bees. Proceedings of the Royal Society B: Biological Sciences 275:797–802. doi:10.1098/rspb.2007.1176

Lawson DA, Whitney HM, Rands SA. 2017. Colour as a backup for scent in the presence of olfactory noise: testing the efficacy backup hypothesis using bumblebees (Bombus terrestris). Royal Society Open Science 4:170996. doi:10.1098/rsos.170996

Lenth R. 2018. emmeans: Estimated Marginal Means, aka Least-Squares Means.

Lotto RB, Chittka L. 2005. Seeing the light: Illumination as a contextual cue to color choice behavior in bumblebees. Proc Natl Acad Sci U S A 102:3852–3856. doi:10.1073/pnas.0500681102

Magnusson A, Skaug H, Nielsen A, Berg C, Kristensen K, Maechler M, Bentham K van, Bolker B, Sadat N, Lüdecke D, Lenth R, O’Brien J, Brooks M. 2020. glmmTMB: Generalized Linear Mixed Models using Template Model Builder.

Marden JH. 1984. Remote perception of floral nectar by bumblebees. Oecologia 64:232–240. doi:10.1007/BF00376876

Menzel R, Blakers M. 1976. Colour receptors in the bee eye—morphology and spectral sensitivity. Journal of comparative physiology 108:11–13.

Menzel R, Giurfa M. 2001. Cognitive architecture of a mini-brain: the honeybee. Trends in Cognitive Sciences 5:62–71. doi:10.1016/S1364-6613(00)01601-6

Miyake T, Yamaoka R, Yahara T. 1998. Floral scents of hawkmoth-pollinated flowers in Japan. J Plant Res 111:199–205. doi:10.1007/BF02512170

Mlinarić A, Horvat M, Šupak Smolčić V. 2017. Dealing with the positive publication bias: Why you should really publish your negative results. Biochemia Medica 27:447–452. doi:10.11613/BM.2017.030201

Muszynski NM, Couvillon PA. 2015. Relational learning in honeybees (Apis mellifera): Oddity and nonoddity discrimination. Behavioural Processes 115:81–93. doi:10.1016/j.beproc.2015.03.001

Oliphant TE. 2006. A guide to NumPy. Trelgol Publishing USA.

R Core Team. 2020. R: A Language and Environment for Statistical Computing. Vienna, Austria: R Foundation for Statistical Computing.

Rering CC, Beck JJ, Hall GW, McCartney MM, Vannette RL. 2018. Nectar-inhabiting microorganisms influence nectar volatile composition and attractiveness to a generalist pollinator. New Phytologist 220:750–759. doi:10.1111/nph.14809

Rhinehart-Doty JA, Schumm J, Smith JC, Smith GP. 1994. A non-taste cue of sucrose in short-term taste tests in rats. Chemical Senses 19:425–431. doi:10.1093/chemse/19.5.425

Rößler DC, Kim K, De Agrò M, Jordan A, Galizia CG, Shamble PS. 2022. Regularly occurring bouts of retinal movements suggest an REM sleep–like state in jumping spiders. Proceedings of the National Academy of Sciences 119:e2204754119. doi:10.1073/pnas.2204754119

Ruck P. 1958. A comparison of the electrical responses of compound eyes and dorsal ocelli in four insect species. Journal of Insect Physiology 2:261–274. doi:10.1016/0022-1910(58)90012-X

Schaeffer RN, Mei YZ, Andicoechea J, Manson JS, Irwin RE. 2017. Consequences of a nectar yeast for pollinator preference and performance. Functional Ecology 31:613–621. doi:10.1111/1365-2435.12762

Sengar M, Narula AK. 2019. Luminescence Sensitization of Eu(III) Complexes with Aromatic Schiff Base and N,N’-Donor Heterocyclic Ligands: Synthesis, Luminescent Properties and Energy Transfer. J Fluoresc 29:111–120. doi:10.1007/s10895-018-2315-3

Skorupski P, Döring TF, Chittka L. 2007. Photoreceptor spectral sensitivity in island and mainland populations of the bumblebee, Bombus terrestris. Journal of Comparative Physiology A 193:485–494.

Srinivasan MV. 2010. Honey Bees as a Model for Vision, Perception, and Cognition. Annual Review of Entomology 55:267–284. doi:10.1146/annurev.ento.010908.164537

Ushey K, Allaire JJ, Tang Y. 2021. reticulate: Interface to “Python.”

van der Walt S, Colbert SC, Varoquaux G. 2011. The NumPy Array: A Structure for Efficient Numerical Computation. Comput Sci Eng 13:22–30. doi:10.1109/MCSE.2011.37

Van Rossum G, Drake FL. 2009. Python 3 Reference Manual. Scotts Valley, CA: CreateSpace.

Von Arx M, Goyret J, Davidowitz G, Raguso RA. 2012. Floral humidity as a reliable sensory cue for profitability assessment by nectar-foraging hawkmoths. Proceedings of the National Academy of Sciences 109:9471–9476.

Waskom M, Botvinnik O, O’Kane D, Hobson P, Lukauskas S, Gemperline DC, Augspurger T, Halchenko Y, Cole JB, Warmenhoven J, Ruiter J de, Pye C, Hoyer S, Vanderplas J, Villalba S, Kunter G, Quintero E, Bachant P, Martin M, Meyer K, Miles A, Ram Y, Yarkoni T, Williams ML, Evans C, Fitzgerald C, Brian Fonnesbeck C, Lee A, Qalieh A. 2017. mwaskom/seaborn: v0.8.1 (September 2017). doi:10.5281/zenodo.883859

Werner A, Menzel R, Wehrhahn C. 1988. Color constancy in the honeybee. Journal of Neuroscience 8:156–159.

Zhang HP, Be’er A, Florin E-L, Swinney HL. 2010. Collective motion and density fluctuations in bacterial colonies. PNAS 107:13626–13630. doi:10.1073/pnas.1001651107

