## Supplementary material for "Bees differentiate sucrose solution from water at a distance": SI - analysis script: S2_Analysis.html

ESM2 Analysis


### ESM2 Analysis

Abstract

This supplement provides the entire R script and output of the
statistical analysis we performed and figures produced, in their
original form. It is presented in the spirit of open and transparent
science, but has not been carefully curated.

### Setup

#### Prepare R environment

```
library(readODS) #to read raw data
library(glmmTMB) #for mixed models
library(car) #for anova on mixed models
library(DHARMa) #for goodness of fit of the model
library(emmeans) #for post hoc
library(ggplot2) #to plot
library(reticulate)

use_python('/home/massimodeagro/anaconda3/envs/DataAnalysis/bin/python')
```

#### Prepare Python environment

```
import pandas as pd
import os
import matplotlib.pyplot as plt
import numpy as np
import seaborn as sns
```

#### Load data

Will load as separate data frames the three conditions and the
spectral data

```
drops <- read.csv(paste0(path,'cond1_Drops.csv'))
tubes <- read.csv(paste0(path,'cond2_Tubes.csv'))
elevated <- read.csv(paste0(path,'cond3_Elevated.csv'))

LightSpectrum <- read.csv(paste0(path,'LightSpectra.csv'))
SolutionSpectrum <- read.csv(paste0(path,'SolutionsSpectra.csv'))

drops$First_Choice <- as.numeric(drops$First_Choice)
drops$Position_Flower <- as.factor(drops$Position_Flower)
drops$Position_Sugar <- as.factor (drops$Position_Sugar)
drops$Position_Water <- as.factor(drops$Position_Water)
drops$Looked_Both <- as.numeric(drops$Looked_Both)
drops$Looked_Both <- as.factor(drops$Looked_Both)
drops$Condition <- "drops"

tubes$First_Choice <- as.numeric(tubes$First_Choice) 
tubes$Position_Flower <- as.factor(tubes$Position_Flower)
tubes$Position_Sugar <- as.factor (tubes$Position_Sugar)
tubes$Position_Water <- as.factor(tubes$Position_Water)
tubes$Looked_Both <- as.numeric(tubes$Looked_Both)
tubes$Looked_Both <- as.factor(tubes$Looked_Both)
tubes$Condition <- "tubes"

elevated$First_Choice <- as.numeric(elevated$First_Choice)
elevated$Position_Flower <- as.factor(elevated$Position_Flower)
elevated$Position_Sugar <- as.factor(elevated$Position_Sugar)
elevated$Position_Water <- as.factor(elevated$Position_Water)
elevated$Looked_Both <- as.numeric(elevated$Looked_Both)
elevated$Looked_Both <- as.factor(elevated$Looked_Both)
elevated$Condition <- "elevated"

elevatedToMerge <- subset(elevated,select = -c(First_Choice_Video,Looked_At_Video,Looked_Both_Video) )
all <- rbind(drops, tubes, elevatedToMerge)
```

```
drops = pd.read_csv(path+'cond1_Drops.csv')
tubes = pd.read_csv(path+'cond2_Tubes.csv')
elevated = pd.read_csv(path+'cond3_Elevated.csv')

LightSpectrum = pd.read_csv(path+'LightSpectra.csv')
SolutionSpectrum = pd.read_csv(path+'SolutionsSpectra.csv')

drops['Condition'] = 'drops'
tubes['Condition'] = 'tubes'
elevated['Condition'] = 'elevated'

elevatedToMerge = elevated
elevatedToMerge.drop(['First_Choice_Video','Looked_At_Video','Looked_Both_Video'], axis=1)
```

```
##          Date Colony  Bee_ID  ... Who_Tested  video_recorded  Condition
## 0    08.10.21   Doris     69  ...     Melina               y   elevated
## 1    08.10.21   Doris     69  ...     Melina               n   elevated
## 2    08.10.21   Doris     69  ...     Melina               y   elevated
## 3    08.10.21   Doris     69  ...     Melina               y   elevated
## 4    08.10.21   Doris     69  ...     Melina               y   elevated
## ..        ...     ...    ...  ...        ...             ...        ...
## 265  10.11.21   Franz     55  ...     Melina               y   elevated
## 266  10.11.21   Franz     55  ...     Melina               y   elevated
## 267  10.11.21   Franz     55  ...     Melina               y   elevated
## 268  10.11.21   Franz     55  ...     Melina               y   elevated
## 269  10.11.21   Franz     55  ...     Melina               y   elevated
## 
## [270 rows x 22 columns]
```

```
all = pd.concat([drops, tubes, elevatedToMerge])
```

### Spectral Analysis

#### Light in the room

We want to plot the spectrum of the two light sources available in
the room. The measure have been taken from under the plexiglas covering
the flight arena, to ensure that the light condition reported is
realistic to the bees experience.

```
# the color matching function has been taken from Dan Burton, see link below.
def w_to_rgb(wavelength, gamma=0.8):

    '''This converts a given wavelength of light to an
    approximate RGB color value. The wavelength must be given
    in nanometers in the range from 380 nm through 750 nm
    (789 THz through 400 THz).

    Based on code by Dan Bruton
    http://www.physics.sfasu.edu/astro/color/spectra.html
    '''

    wavelength = float(wavelength)
    if wavelength >= 380 and wavelength <= 440:
        attenuation = 0.3 + 0.7 * (wavelength - 380) / (440 - 380)
        R = ((-(wavelength - 440) / (440 - 380)) * attenuation) ** gamma
        G = 0.0
        B = (1.0 * attenuation) ** gamma
    elif wavelength >= 440 and wavelength <= 490:
        R = 0.0
        G = ((wavelength - 440) / (490 - 440)) ** gamma
        B = 1.0
    elif wavelength >= 490 and wavelength <= 510:
        R = 0.0
        G = 1.0
        B = (-(wavelength - 510) / (510 - 490)) ** gamma
    elif wavelength >= 510 and wavelength <= 580:
        R = ((wavelength - 510) / (580 - 510)) ** gamma
        G = 1.0
        B = 0.0
    elif wavelength >= 580 and wavelength <= 645:
        R = 1.0
        G = (-(wavelength - 645) / (645 - 580)) ** gamma
        B = 0.0
    elif wavelength >= 645 and wavelength <= 750:
        attenuation = 0.3 + 0.7 * (750 - wavelength) / (750 - 645)
        R = (1.0 * attenuation) ** gamma
        G = 0.0
        B = 0.0
    else:
        R = 0.0
        G = 0.0
        B = 0.0

    #R *= 255
    #G *= 255
    #B *= 255
    return (R, G, B)

all_colors = []
for row in range(len(LightSpectrum)):
    current_wavelength = LightSpectrum['Wavelength(nm)'][row]

    all_colors.append(w_to_rgb(current_wavelength))

for i in range(len(LightSpectrum)-1):
    plt.axvspan(LightSpectrum['Wavelength(nm)'][i],LightSpectrum['Wavelength(nm)'][i+1], color=all_colors[i])

plt.plot(LightSpectrum['Wavelength(nm)'],LightSpectrum['LED'],color="grey")
plt.plot(LightSpectrum['Wavelength(nm)'],LightSpectrum['Neon'],color="white")
plt.axvline(300, c='#aaaaaa', lw=3, ls='--')
plt.axvline(600, c='#aaaaaa', lw=3, ls='--')
plt.show()
```

```
plt.close()
```

this one is fore completeness. The plot that will go in the paper
only includes bee visual spectrum

```
for i in range(len(LightSpectrum)-1):
    plt.axvspan(LightSpectrum['Wavelength(nm)'][i],LightSpectrum['Wavelength(nm)'][i+1], color=all_colors[i])

plt.plot(LightSpectrum['Wavelength(nm)'],LightSpectrum['LED'],color="grey")
plt.plot(LightSpectrum['Wavelength(nm)'],LightSpectrum['Neon'],color="white")

plt.xlim(300, 600)
```

```
## (300.0, 600.0)
```

```
plt.show()
```

```
plt.close()
```

we seem to not have uv components, with no light until ~400nm.

#### Drops appearance

We will plot transmittance and reflectance of both the sucrose
solution and water. We won’t plot transmittance as can be easily derived
from absorbance, and would give a very similar information

```
from matplotlib.lines import Line2D
custom_lines = [Line2D([0], [0], color='#ffffff', lw=4),
                Line2D([0], [0], color='#999999', lw=4)]

all_colors = []
for row in range(len(SolutionSpectrum)):
    current_wavelength = SolutionSpectrum['Wavelength'][row]

    all_colors.append(w_to_rgb(current_wavelength))

fig, axs = plt.subplots(2,1)
for i in range(len(SolutionSpectrum)-1):
    axs[0].axvspan(SolutionSpectrum['Wavelength'][i],SolutionSpectrum['Wavelength'][i+1], color=all_colors[i])
axs[0].plot(SolutionSpectrum['Wavelength'].values, SolutionSpectrum['tap_water_Absorbance'].values, c='#ffffff')
axs[0].plot(SolutionSpectrum['Wavelength'].values, SolutionSpectrum['tap_sugar_Absorbance'].values, c='#999999')
axs[0].set_xlim(300,600)
```

```
## (300.0, 600.0)
```

```
for i in range(len(SolutionSpectrum)-1):
    axs[1].axvspan(SolutionSpectrum['Wavelength'][i],SolutionSpectrum['Wavelength'][i+1], color=all_colors[i])
axs[1].plot(SolutionSpectrum['Wavelength'].values, SolutionSpectrum['tap_water_Reflectance'].values, c='#ffffff')
axs[1].plot(SolutionSpectrum['Wavelength'].values, SolutionSpectrum['tap_sugar_Reflectance'].values, c='#999999')
axs[1].legend(custom_lines, ['Water', 'Sucrose'], loc='lower right')
axs[1].set_xlim(300,600)
```

```
## (300.0, 600.0)
```

```
plt.show()
```

```
plt.close()
```

These two graphs anyway suggest that the two solutions should be
discernible by the bees

### Data Analysis

#### Preliminary Analysis - effect of reward position

Before proceeding with the main analysis, we want to see if the bees
had any type of bias, in term of the side of the sucrose solution in the
flower, and the position (1-6)

```
mp0 <- glmmTMB(First_Choice~Position_Flower*Position_Sugar*Condition+(1|Bee_ID_Long), family = binomial, data = all)

simres <- simulateResiduals(mp0)
plot(simres)
```

model fit is good. we proceed

```
Anova(mp0)
```

```
## Analysis of Deviance Table (Type II Wald chisquare tests)
## 
## Response: First_Choice
##                                            Chisq Df Pr(>Chisq)    
## Position_Flower                          13.0830  5  0.0226130 *  
## Position_Sugar                           20.1606  3  0.0001572 ***
## Condition                                 3.1588  2  0.2060993    
## Position_Flower:Position_Sugar           45.5712 15  6.219e-05 ***
## Position_Flower:Condition                 3.3243 10  0.9727296    
## Position_Sugar:Condition                  5.6127  6  0.4679453    
## Position_Flower:Position_Sugar:Condition 31.7191 30  0.3807039    
## ---
## Signif. codes:  0 '***' 0.001 '**' 0.01 '*' 0.05 '.' 0.1 ' ' 1
```

We found an overall effect of both flower position and the side in
which the sugar is, while we found no interaction with experimental
condition. This seems to imply that bees were biased toward some
combination, but this had no effect on the treatment, just a
pre-existing preference. Will proceed with a post-hoc

```
e<- emmeans(mp0, ~Position_Flower+Position_Sugar, type='response')
```

```
## NOTE: Results may be misleading due to involvement in interactions
```

```
e
```

```
##  Position_Flower Position_Sugar  prob     SE  df lower.CL upper.CL
##  1               b              0.515 0.0811 721    0.360    0.668
##  2               b              0.413 0.0892 721    0.255    0.592
##  3               b              0.447 0.0903 721    0.283    0.623
##  4               b              0.660 0.0892 721    0.471    0.809
##  5               b              0.620 0.0952 721    0.424    0.783
##  6               b              0.325 0.0999 721    0.164    0.540
##  1               f              0.663 0.0745 721    0.506    0.791
##  2               f              0.626 0.1154 721    0.388    0.815
##  3               f              0.777 0.0658 721    0.623    0.880
##  4               f              0.635 0.1043 721    0.418    0.808
##  5               f              0.731 0.0950 721    0.513    0.875
##  6               f              0.841 0.0767 721    0.632    0.942
##  1               l              0.332 0.0824 721    0.193    0.507
##  2               l              0.203 0.0551 721    0.115    0.332
##  3               l              0.329 0.1080 721    0.158    0.561
##  4               l              0.707 0.0961 721    0.492    0.857
##  5               l              0.810 0.0639 721    0.653    0.906
##  6               l              0.558 0.1007 721    0.361    0.737
##  1               r              0.814 0.0731 721    0.629    0.919
##  2               r              0.614 0.0830 721    0.444    0.759
##  3               r              0.808 0.0774 721    0.613    0.918
##  4               r              0.468 0.1208 721    0.253    0.695
##  5               r              0.413 0.1033 721    0.234    0.619
##  6               r              0.628 0.1003 721    0.421    0.797
## 
## Results are averaged over the levels of: Condition 
## Confidence level used: 0.95 
## Intervals are back-transformed from the logit scale
```

```
ggplot(as.data.frame(e), aes(x=Position_Flower, y=prob, color=Position_Sugar))+
  geom_point(position = position_dodge(0.8))+
  geom_errorbar(aes(ymin=prob-SE, ymax=prob+SE),position = position_dodge(0.8))+
  ylim(0,1)
```

Testing all the pairs seems superfluous. It seems in general that
bees prefer drops that are towards the center of the arena, rather than
towards the wall: - for positions 1, 2, and 3, which are near the left
wall, drops on the right are preferred - for position 4, 5, and 6, near
the right wall, drops on the left are preferred There is also an overall
preference for drops towards the front rather than the back, which is
easy to explain being the first encountered between the two in each
visit.

#### Preliminary Analysis - time

as a second preliminary measure, we want to know whether the time
spent from flight arena entrance to first choice, across conditions and
across trials. This should represent a measure of procedure learning
speed.

```
mp1 <- glmmTMB(Time_To_Touch_Flower..s.~Bout_Number*Condition+(1|Bee_ID_Long), family = gaussian, data = all) #TODO fix model fit

simres <- simulateResiduals(mp1)
plot(simres)
```

model fit is good. we proceed

```
Anova(mp1)
```

```
## Analysis of Deviance Table (Type II Wald chisquare tests)
## 
## Response: Time_To_Touch_Flower..s.
##                         Chisq Df Pr(>Chisq)    
## Bout_Number           54.5895  1  1.485e-13 ***
## Condition              8.4043  2   0.014964 *  
## Bout_Number:Condition  9.7264  2   0.007726 ** 
## ---
## Signif. codes:  0 '***' 0.001 '**' 0.01 '*' 0.05 '.' 0.1 ' ' 1
```

there are effects related to condition, that probably have to do with
the learning procedure. The crucial effect is with bout number and
interaction, as it indeed suggests a learning speed if the value is
decreasing.

```
et <- emtrends(mp1, ~Condition, var='Bout_Number', type='response')
test(et, adjust='bonferroni')
```

```
##  Condition Bout_Number.trend    SE  df t.ratio p.value
##  drops                 -3.02 0.719 786  -4.203  0.0001
##  elevated              -1.48 0.706 786  -2.101  0.1078
##  tubes                 -4.60 0.708 786  -6.499  <.0001
## 
## P value adjustment: bonferroni method for 3 tests
```

time to touch the reward decreases both in drops and tubes, but not
in elevated.

##### sub-check. is time a good predictor for look at both?

we measured a dicotomical variable: looked at both. this signifies
that the bees inspected both drops before selecting one. Expecially in
further trials, is the time taken before decision a good predictor of
this behaviour? We will keep in bout number as it is a clear
moderator.

```
mp2 <- glmmTMB(Time_To_Touch_Flower..s.~Looked_Both*Bout_Number+(1|Bee_ID_Long), family = gaussian, data = all) #TODO fix model fit

simres <- simulateResiduals(mp2)
plot(simres)
```

```
Anova(mp2)
```

```
## Analysis of Deviance Table (Type II Wald chisquare tests)
## 
## Response: Time_To_Touch_Flower..s.
##                           Chisq Df Pr(>Chisq)    
## Looked_Both             12.1913  1  0.0004801 ***
## Bout_Number             55.6808  1  8.525e-14 ***
## Looked_Both:Bout_Number  5.7775  1  0.0162322 *  
## ---
## Signif. codes:  0 '***' 0.001 '**' 0.01 '*' 0.05 '.' 0.1 ' ' 1
```

```
e <- emmeans(mp2, ~Looked_Both)
```

```
## NOTE: Results may be misleading due to involvement in interactions
```

```
e
```

```
##  Looked_Both emmean   SE  df lower.CL upper.CL
##  0             9.75 1.80 788     6.21     13.3
##  1            17.42 1.77 788    13.94     20.9
## 
## Confidence level used: 0.95
```

```
pairs(e)
```

```
##  contrast estimate   SE  df t.ratio p.value
##  0 - 1       -7.67 2.19 788  -3.498  0.0005
```

```
ggplot(all, aes(x=Bout_Number, y=Time_To_Touch_Flower..s., color=Looked_Both))+
  facet_wrap(~Condition)+
        geom_point()+
        geom_smooth(method='glm')
```

```
## `geom_smooth()` using formula 'y ~ x'
```

```
## Warning: Removed 16 rows containing non-finite values (stat_smooth).
```

```
## Warning: Removed 16 rows containing missing values (geom_point).
```

This is very interesting. some points about this: - Elevated
condition have very short times. This seems to be a very evident
condition, that brings bees to immediately locate drops. - the other
conditions have an initial longer time, that gets low towards the end of
the experiment. When the animal didn’t inspect both instead, the time
remain very low. - overall, it seems that visit time is a good predictor
for looked at both

#### Main analysis

to the main observation. Are the bumblebees able to discriminate
sugar and water? in which condition? Is this dependent on visit number
and their willingness to look at both drops?

```
m0 <- glmmTMB (First_Choice~Looked_Both* Bout_Number* Condition + (1|Bee_ID_Long), data=all, family=binomial)
simres <- simulateResiduals(m0)
plot(simres)
```

```
Anova(m0)
```

```
## Analysis of Deviance Table (Type II Wald chisquare tests)
## 
## Response: First_Choice
##                                     Chisq Df Pr(>Chisq)    
## Looked_Both                       39.4480  1  3.369e-10 ***
## Bout_Number                        9.4473  1   0.002115 ** 
## Condition                          2.7484  2   0.253038    
## Looked_Both:Bout_Number            2.0519  1   0.152021    
## Looked_Both:Condition              0.1360  2   0.934260    
## Bout_Number:Condition              1.4637  2   0.481009    
## Looked_Both:Bout_Number:Condition  0.1969  2   0.906225    
## ---
## Signif. codes:  0 '***' 0.001 '**' 0.01 '*' 0.05 '.' 0.1 ' ' 1
```

- There is a big effect of looked at both. Probably due to the bees
  being better at discriminating when they take the time to actually
  inspect.
- There is an effect of bout number, probably a component of
  learning
- No difference between conditions (however we are gonna keep the
  predictor, as they have been conducted as separate experiments and would
  be disingenuous to pool all data together

```
et <- emtrends(m0, ~Condition*Looked_Both, var='Bout_Number', type='response')
test(et, adjust='bonferroni')
```

```
##  Condition Looked_Both Bout_Number.trend     SE  df t.ratio p.value
##  drops     0                      0.0800 0.0732 781   1.093  1.0000
##  elevated  0                      0.0807 0.0623 781   1.295  1.0000
##  tubes     0                     -0.0161 0.0805 781  -0.201  1.0000
##  drops     1                      0.1296 0.0736 781   1.762  0.4711
##  elevated  1                      0.1776 0.0860 781   2.065  0.2355
##  tubes     1                      0.0956 0.0633 781   1.510  0.7893
## 
## P value adjustment: bonferroni method for 6 tests
```

The trend seem all to be of overall increase in correct responses,
excluded tubes looked at only one drop that is just above 1% per bout.
None however result significant, probably due to the correction.
Interesting regardless. Now to the main result.

```
e <- emmeans(m0, ~Condition*Looked_Both, type='response')
```

```
## NOTE: Results may be misleading due to involvement in interactions
```

```
test(e, adjust='bonferroni')
```

```
##  Condition Looked_Both  prob     SE  df null t.ratio p.value
##  drops     0           0.490 0.0480 781  0.5  -0.204  1.0000
##  elevated  0           0.487 0.0400 781  0.5  -0.313  1.0000
##  tubes     0           0.415 0.0454 781  0.5  -1.827  0.4084
##  drops     1           0.721 0.0373 781  0.5   5.118  <.0001
##  elevated  1           0.698 0.0460 781  0.5   3.836  0.0008
##  tubes     1           0.663 0.0394 781  0.5   3.838  0.0008
## 
## P value adjustment: bonferroni method for 6 tests 
## Tests are performed on the logit scale
```

```
contrast(e, list('Drops'=c(0.5,0,0,0.5,0,0),
                 'Tubes'=c(0,0,0.5,0,0,0.5),
                 'Elevated'=c(0,0.5,0,0,0.5,0),
                'Drops_InspectedBoth' = c(0,0,0,1,0,0),
                'Drops_InspectedOne' = c(1,0,0,0,0,0),
                'Tubes_InspectedBoth' = c(0,0,0,0,0,1),
                'Tubes_InspectedOne' = c(0,0,1,0,0,0),
                'Elevated_InspectedBoth' = c(0,0,0,0,1,0),
                'Elevated_InspectedOne' = c(0,1,0,0,0,0)), adjust='bonferroni')
```

```
## Note: Use 'contrast(regrid(object), ...)' to obtain contrasts of back-transformed estimates
```

```
##  contrast               estimate    SE  df t.ratio p.value
##  Drops                    0.4549 0.133 781   3.410  0.0062
##  Tubes                    0.1675 0.128 781   1.304  1.0000
##  Elevated                 0.3933 0.135 781   2.907  0.0338
##  Drops_InspectedBoth      0.9489 0.185 781   5.118  <.0001
##  Drops_InspectedOne      -0.0391 0.192 781  -0.204  1.0000
##  Tubes_InspectedBoth      0.6765 0.176 781   3.838  0.0012
##  Tubes_InspectedOne      -0.3415 0.187 781  -1.827  0.6125
##  Elevated_InspectedBoth   0.8368 0.218 781   3.836  0.0012
##  Elevated_InspectedOne   -0.0501 0.160 781  -0.313  1.0000
## 
## Note: contrasts are still on the logit scale 
## P value adjustment: bonferroni method for 9 tests
```

When the bees inspected both drops, they were able to discriminate
sucrose from water in all conditions, while when they did not inspect,
the choice dropped to random. When pooling all together, only drop and
Elevated condition remains significant, while tubes drops to random.
This is probably to do with the task difficulty and low visibility of
the drops in this condition. Indeed, when not looking both in tube
condition they don’t seem to get better over time, signifying that
inspection is really important to get the relevant info here.

```
g = sns.FacetGrid(all, col="Condition", hue="Looked_Both")
g.map(sns.regplot, "Bout_Number", "First_Choice", x_jitter=.3, y_jitter=.03, scatter_kws={'s':5})
```

```
g.add_legend()
```

#### Differences between manual and video analysis

we collected videos for the third condition of our experiment, to
make sure our manual scoring was overall unbiased. In this section, we
will compare the data obtained manually and with video from a second
experimenter, to see the level of internal consistency

```
elevated <- read.csv(paste0(path,'cond3_Elevated.csv')) #reloading to set as numeric

cor.test(elevated$Looked_Both, elevated$Looked_Both_Video)
```

```
## 
##  Pearson's product-moment correlation
## 
## data:  elevated$Looked_Both and elevated$Looked_Both_Video
## t = 11.322, df = 253, p-value < 2.2e-16
## alternative hypothesis: true correlation is not equal to 0
## 95 percent confidence interval:
##  0.4920998 0.6560002
## sample estimates:
##       cor 
## 0.5798877
```

```
cor.test(elevated$First_Choice, elevated$First_Choice_Video)
```

```
## 
##  Pearson's product-moment correlation
## 
## data:  elevated$First_Choice and elevated$First_Choice_Video
## t = 30.779, df = 253, p-value < 2.2e-16
## alternative hypothesis: true correlation is not equal to 0
## 95 percent confidence interval:
##  0.8593180 0.9117269
## sample estimates:
##       cor 
## 0.8883822
```

```
sum(elevated$Looked_Both, na.rm = TRUE)
```

```
## [1] 106
```

```
mean(elevated$Looked_Both, na.rm = TRUE)
```

```
## [1] 0.3984962
```

```
sum(elevated$Looked_Both_Video, na.rm = TRUE)
```

```
## [1] 70
```

```
mean(elevated$Looked_Both_Video, na.rm = TRUE)
```

```
## [1] 0.2702703
```

```
elevated_onlylookvideo <- subset(elevated, elevated$Looked_Both_Video == 1)

mean(elevated_onlylookvideo$Looked_Both_Video, na.rm = TRUE)
```

```
## [1] 1
```

```
mean(elevated_onlylookvideo$Looked_Both, na.rm = TRUE)
```

```
## [1] 0.8571429
```

First choice is congruent 88.8% of the time, revealing a high
internal consistency. The difference mostly depends on some trial in
which bees tasted the sugar rapidly before landing, which was not
noticed during manual scoring and as such not included. For the same
reason, most of these occurrences were initially scored as “looked at
both” as the bee hovered in front of the tasted in flight drop before
choosing the other. Completely different story the look at both
condition, being congruent only 57.9% of the times. This suggests that
the measure is a bit too dependent on the scorer perception, and as such
we hereby decide against using it for making claims.

We will proceed with the same analysis now, to check whether the
results tell the same story even after video analysis

```
mc0 <- glmmTMB (First_Choice_Video~Looked_Both_Video* Bout_Number + (1|Bee_ID_Long), data=elevated, family=binomial)
simres <- simulateResiduals(mc0)
plot(simres)
```

```
Anova(mc0)
```

```
## Analysis of Deviance Table (Type II Wald chisquare tests)
## 
## Response: First_Choice_Video
##                                Chisq Df Pr(>Chisq)   
## Looked_Both_Video             7.4756  1   0.006254 **
## Bout_Number                   5.8149  1   0.015891 * 
## Looked_Both_Video:Bout_Number 1.6081  1   0.204766   
## ---
## Signif. codes:  0 '***' 0.001 '**' 0.01 '*' 0.05 '.' 0.1 ' ' 1
```

```
et <- emtrends(mc0, ~Looked_Both_Video, var='Bout_Number', type='response')
test(et, adjust='bonferroni')
```

```
##  Looked_Both_Video Bout_Number.trend    SE  df t.ratio p.value
##                  0            0.0934 0.057 254   1.640  0.2046
##                  1            0.2627 0.121 254   2.176  0.0610
## 
## P value adjustment: bonferroni method for 2 tests
```

```
e <- emmeans(mc0, ~Looked_Both_Video, type='response')
```

```
## NOTE: Results may be misleading due to involvement in interactions
```

```
e
```

```
##  Looked_Both_Video  prob     SE  df lower.CL upper.CL
##                  0 0.540 0.0367 254    0.468    0.611
##                  1 0.746 0.0550 254    0.623    0.838
## 
## Confidence level used: 0.95 
## Intervals are back-transformed from the logit scale
```

```
contrast(e, list('looked_one'=c(1,0),
                 'looked_both'=c(0,1),
                 'together'=c(0.5,0.5)), adjust='bonferroni')
```

```
## Note: Use 'contrast(regrid(object), ...)' to obtain contrasts of back-transformed estimates
```

```
##  contrast    estimate    SE  df t.ratio p.value
##  looked_one     0.161 0.148 254   1.093  0.8263
##  looked_both    1.075 0.290 254   3.707  0.0008
##  together       0.618 0.163 254   3.799  0.0005
## 
## Note: contrasts are still on the logit scale 
## P value adjustment: bonferroni method for 3 tests
```

bees chose sugar 54% of the times when didn’t inspect both, 74.6%
when they did. overall, they chose sugar 64.3% of the times. the result
look identical to the manually scored ones. even the inspection
variable, in the end appears consistent. the difference must be equally
distributed: same number of formerly inspected both being scored as
inspected one, in respect to formerly inspected one being scored as
inspected both.
